# High-THC *Cannabis* smoke impairs working memory capacity in spontaneous tests of novelty preference for objects and odors in rats

**DOI:** 10.1101/2023.04.06.535880

**Authors:** Ilne L. Barnard, Timothy J. Onofrychuk, Aaron D. Toderash, Vyom N. Patel, Aiden E. Glass, Jesse C. Adrian, Robert. B. Laprairie, John G. Howland

## Abstract

Working memory (WM) is an executive function that orchestrates the use of a limited amount of information, referred to as working memory capacity (WMC), in cognitive functions. In humans, *Cannabis* exposure impairs WM; however, it is unclear if *Cannabis* facilitates or impairs rodent WM. Existing literature also fails to address the effects of *Cannabis* exposure on rodent WMC using exposure paradigms that closely mirror patterns of human use. In the present study, WMC of rats was inferred by novelty preference after a short delay in spontaneous recognition-based tests. Either object or odor-based stimuli were used in different variations of the tests that present identical (IOT) and different (DOT) sets of stimuli (3 or 6) for low-and high-cognitive loads, respectively. Additionally, we present a human-machine hybrid (HYB) behavioral quantification approach which supplements stopwatch-based scoring with supervised machine learning (SML)-based classification, enabling behavioral data to be made publicly available. After validating the spontaneous tests, 6-item IOT and DOT tests with the HYB method were used to evaluate the impact of acute exposure to high-THC or high-CBD *Cannabis* smoke on novelty preference. Under control conditions, rats showed novelty preference in all test variations. We found that high-THC, but not high-CBD, *Cannabis* smoke exposure impaired novelty preference for objects under a high-cognitive load. Odor-based recognition deficits were seen under both low-, and high-cognitive loads only following high-THC smoke exposure. Ultimately, these data show that *Cannabis* smoke exposure impacts novelty preference in a load-dependent, and stimuli-specific manner.

**Significance Statement:** Working memory (WM) capacity is the limited amount of information that can be utilized by WM to orchestrate processes like learning and memory. Using object-and odor-based spontaneous recognition tests, the impact of high-THC or high-CBD *Cannabis* smoke on novelty preference was evaluated. Behavioral measurements were generated using a combination of open-source analysis software and traditional stopwatch scoring to form a human-machine hybrid (HYB) scoring method. We show novelty preference deficits under high-cognitive loads in object-based tests, while impacting novelty preference under both high-and low-cognitive loads in the odor-based tests. Ultimately, we show that *Cannabis* smoke exposure affects cognitive functions that underly WM in rats, which has broad implications for human use.

## Introduction

Working memory (WM) is an executive function that orchestrates the use of a limited amount of information in cognitive functions like learning and memory (Constantinidis & Klingberg, 2016; D’Esposito et al., 1995; Eriksson et al., 2015; Wilhelm et al., 2013). In humans, Δ^9^-tetrahydrocannabinol (THC), the main psychoactive constituent of *Cannabis*, impairs WM following both acute and chronic *Cannabis* exposure, likely by action at the cannabinoid type 1 receptor (CB1R) (Adam et al., 2020; Bossong et al., 2012; Cousijn et al., 2014; Crane et al., 2013; Curran et al., 2002; D’Souza et al., 2012; Ilan et al., 2004; Ligresti et al., 2016; Owens et al., 2019). The WM impairments produced by *Cannabis* have been interpreted as a combination of disruptions in active maintenance, limited capacity, interference control, and flexible updating subconstructs of WM (Barch & Smith, 2008). In contrast, studies in rodents demonstrate both THC-mediated impairments and improvements in WM function (Barnard et al., 2022; Blaes et al., 2019; Bruijnzeel et al., 2016; de Melo et al., 2005; Goonawardena et al., 2010; Varvel et al., 2001). These inconsistent findings may be attributable to differences in the behavioral tasks used, cannabinoid dosage, exposure timelines, and routes of administration (Baglot et al., 2021; Hložek et al., 2017; Klausner & Dingell, 1971; Nguyen et al., 2016; Wiley et al., 2021). Importantly, previous rodent studies have not directly assessed the effects of *Cannabis* exposure on WMC. WMC is essential for higher cognitive operations critical to everyday function and can be impaired in disorders like schizophrenia and Parkinson’s disease (Goldman-Rakic, 1999; Piskulic et al., 2007; Gold et al., 2018).

A shortcoming in rodent literature is that traditional rodent WMC tests mimic n-back or recall WM tests used in humans that require a long training period, learned rules, and considerable experimental involvement (Barnard et al., 2022; Cowan, 2010; Daneman & Carpenter, 1980; Dudchenko, 2004; Dudchenko et al., 2013; Kirchner, 1958; Oomen et al., 2013; Scott et al., 2020; Vorhees & Williams, 2014; Wilhelm et al., 2013). Spontaneous recognition tests circumvent these weaknesses by relying on rodents’ innate novelty seeking behavior as shown by preferential interaction with a novel stimulus after a delay (Broadbent et al., 2004; Ennaceur & Aggleton, 1994; Ennaceur & Delacour, 1988; Sannino et al., 2012). Novelty preference has been used to assess WMC in mice under low-and high-cognitive loads through the Identical Objects Task (IOT) and the Different Objects Task (DOT), respectively (Olivito et al., 2016, 2019; Sannino et al., 2012) Therefore, the first goal of the present study was to validate these spontaneous tests using objects in rats. In addition, we developed and validated olfactory versions of these tests to evaluate novelty preference for odors.

For all test variations, novelty preference was inferred by measuring the relative amount of interaction behavior exhibited at novel and previously experienced stimuli after a short delay. Typical approaches to quantifying rodent behavior for spontaneous interaction tests are generally laborious, prone to human subjectivity, and lack objective analysis steps that can be verified and reproduced (Anderson & Perona, 2014). Recent advances in automated behavioral analysis have enabled researchers to obtain a detailed and objective record of a diversity of complex behaviors across species (Cui et al., 2021; Newton et al., 2023; Nilsson et al., 2020; Slivicki et al., 2023; Winters et al., 2022). Here, we automatically quantified interaction events using a supervised machine learning (SML)-based analysis approach, then upon manual inspection of SML predictions, sub-optimal predictions were supplemented by human stopwatch scoring to form a human-machine hybrid (HYB) scoring method (Mathis et al., 2018; Nilsson et al., 2020). By automatically predicting interaction events frame-by-frame, several secondary behavioral measures, including approach latency and interaction bout count, were easily calculated and provide a more complete characterization of novelty preference to infer WMC. To our knowledge, the present study is the first demonstration of SML-based behavioral analysis in the context of a spontaneous interaction-based test.

Using validated spontaneous tasks and the HYB scoring method, our second goal was to assess the effects of *Cannabis* smoke exposure on novelty preference to infer WM and WMC. We tested rats shortly after acute exposure to the smoke of either high-THC or high-CBD-containing *Cannabis* buds using a validated paradigm (Barnard et al., 2022; Roebuck et al., 2022). We found that high-THC, but not high-CBD, smoke impaired performance of the tests in a stimuli-specific manner.

## Materials & Methods

### Subjects

Adult (2-4 months of age) male (n = 92) Long-Evans rats (Charles River Laboratories, Kingston, NY) were pair housed in a vivarium in standard ventilated cages with ad libitum water and food, and a plastic tube for environmental enrichment on a 12-hour light/dark cycle (starting at 0700). For establishment and validation of the object-and odor-based IOT and DOT tests 52 rats were used, and 48 rats were used to evaluate the impact of acute *Cannabis* smoke exposure on novelty preference. Rats were tested at the same time of day between the hours of 0730 and 1800. All procedures followed guidelines from the Canadian Council on Animal Care and were approved by the local animal research ethics board.

### Apparatus and testing materials

Rats were handled in the testing room (3 mins a day for 3 days) and subsequently habituated to the testing apparatus (10 min for 2 days). Rats were tested in a white corrugated plastic box (60 cm x 60 cm x 60 cm) with the stimuli evenly presented between two opposing walls at three positions (see Fig 1; 9 cm from side of box, 21.5 cm apart from each other). Object stimuli were created from a variety of LEGO™ pieces of different sizes and colors with an average size of 7 cm x 10 cm. LEGO™ was chosen to maintain consistency between different object sets. Odor stimuli were created using 250 mL glass canning jars. The jars were filled with sand for stability, and to provide a resting place for a small plastic vile filled half-way with a powered spice (lemon pepper, dill, sage, onion, anise, cloves, ginger, cumin, cocoa, celery salt, coffee, cinnamon, garlic, or oregano). Holes were drilled in the lids of the jars to allow the rats to smell the spices. All items were affixed to the testing apparatus with Velcro™ at one of six positions to prevent them from being displaced during the test.

**Figure 1.**
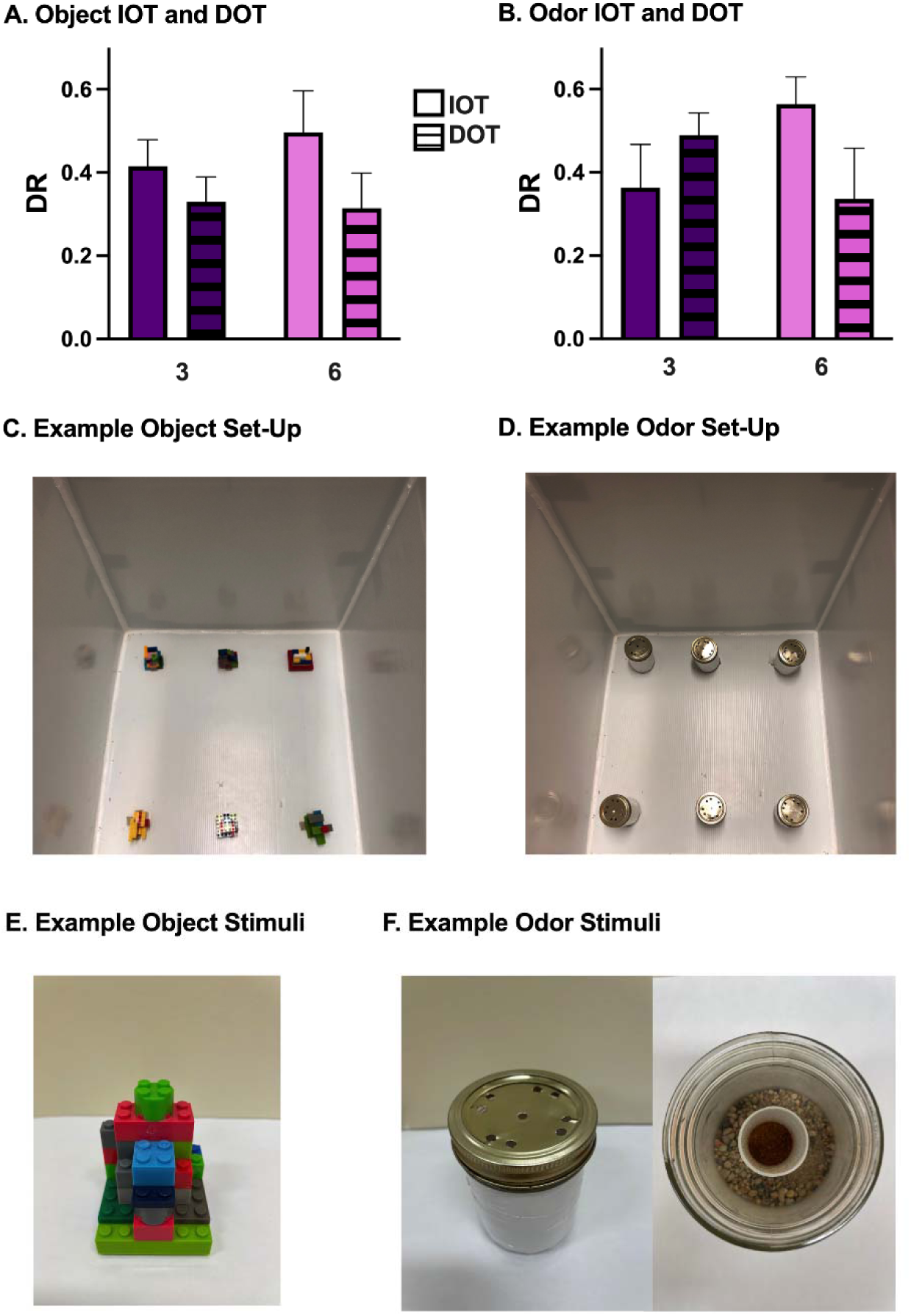
The validation and establishment of the object and odor based IOT and DOT tests. **A** Object interaction was measured using DR’s to evaluate novelty preference using 3- objects and 6-objects. Rats explore the novel object significantly more than the familiar objects in the IOT and DOT variations with both 3- and 6- objects. No differences in novelty preference or exploration times are seen between the IOT and DOT variations, or between 3-object and 6- object versions. **B** Odor interaction was also measured using DR’s to evaluate novelty preference using 3-odors and 6-odors. Rats explore the novel odor significantly more than the familiar odors in the IOT and DOT variations with both 3- and 6- odors. No differences in novelty preference or exploration times are seen between the IOT and DOT variations, or between 3-odor and 6-odor versions. **C** A picture of an example object set-up is shown. Objects are displayed in 6 positions in a white-corrugated plastic box. **D** A picture of an example odor set-up is shown. Odors are displayed in 6 positions in a white-corrugated plastic box. **E** An example of an object stimuli **F** An example of an odor stimuli. Data is represented as mean ± SEM.

### Spontaneous WMC test protocol

To validate the object IOT and DOT, 24 naïve rats performed both the 3- and 6- object variations. Twenty naïve rats were used to establish the 3- and 6-odor IOT and DOT test variations. Using a within-subjects design, 48 rats performed both the IOT and DOT variants either in the object or odor-based tests 20 min after *Cannabis* smoke exposure (see Fig 2). The order of tests was quasi-counterbalanced, and rats had a 2-day washout period between tests. On the test day, the testing box was prepared with 2 sets of 6 stimuli for the test and paradigm being performed (Fig 2). The rat was then placed into the testing box for the sample phase, for a duration of 5 minutes. Following the sample phase, the rat was taken out of the testing box and placed inside a transport cage for 1 min. During the delay, all stimuli were replaced for the test phase. Then, the rat was placed back into the box for the test phase (5 min). The testing box and the stimuli were cleaned with 70% ethanol after each phase.

**Figure 2.**
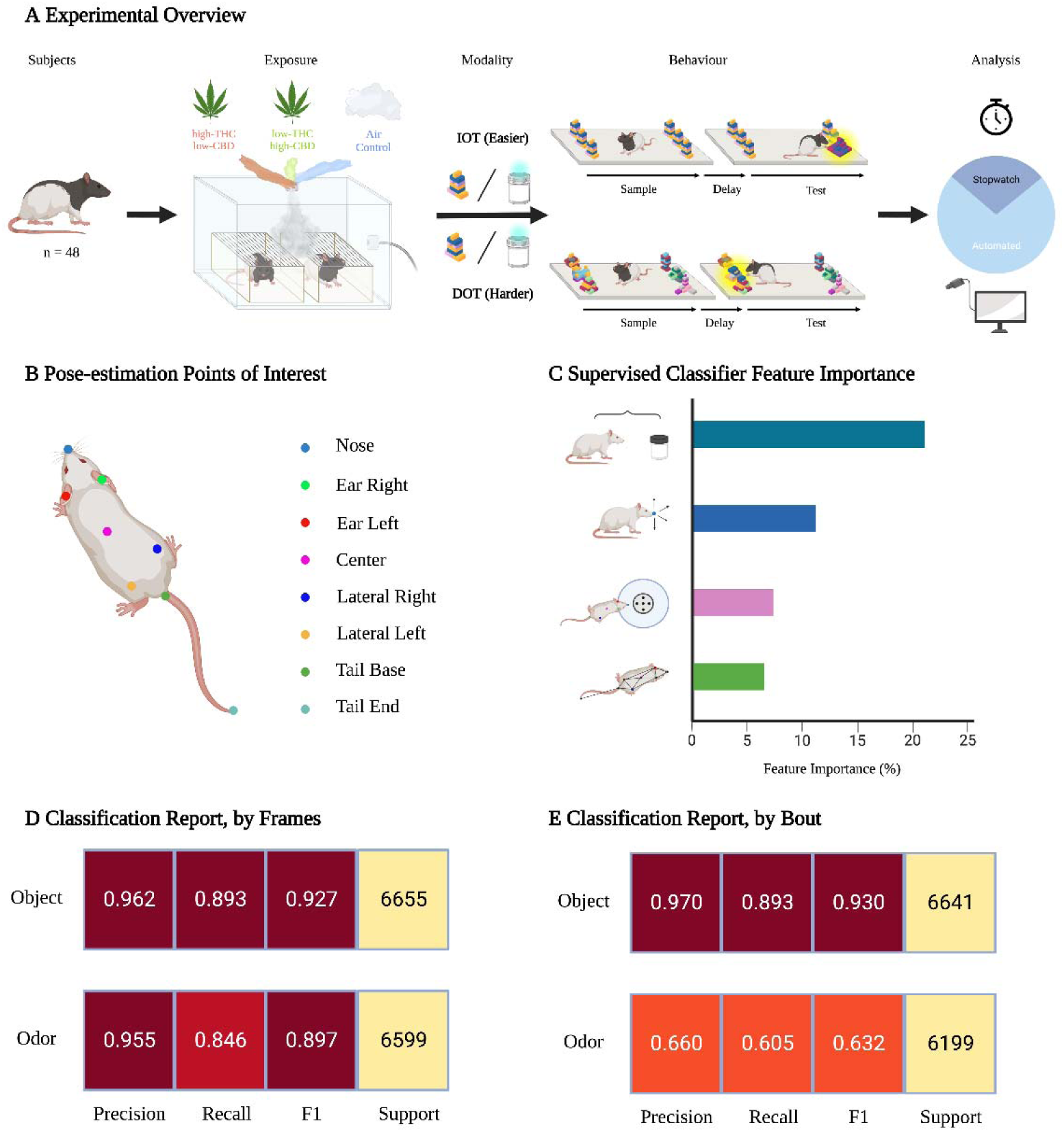
Experimental overview for acute *Cannabis* exposure and behavioral classifier training. **A** Schematic representation of the experimental design. Male Long-Evans rats (n = 48) were used for this study. Using a repeated measures experimental design, each rat was exposed to high-THC *Cannabis* smoke, low-THC *Cannabis* smoke, and an Air Control condition. Rats were exposed 20 minutes prior to the start of behavioral testing. Each rat either underwent the 6- object IOT and 6-object DOT, or the 6-odor IOT and 6-odor DOT. The order in which the IOT and DOT variations were performed was randomized. Rat behavior was quantified using traditional stopwatch scoring and by automated SML-based behavioral analysis. Sub-optimal SML predictions were replaced by stopwatch scoring, constituting a HYB scoring approach. **B** Illustration of the point-of-interest configuration used for pose-estimation analysis. **C** Visualization of the 50 most important features used for model training. Important features were grouped into distance to stimuli, nose movements, region-of-interest, and spatial dynamics between points-of-interest. **D** Classifier performance metrics for the object (top) and odor (bottom) models. Test frames were randomly extracted from the dataset (20% test, 80% train). **E** Classifier performance metrics for the object (top) and odor (bottom) models. Test bouts were randomly extracted from the dataset (20% test, 80% train). This figure was created using BioRender.com.

### Cannabis bud preparation and acute smoke exposure protocol

A high-THC (19.51%) and low-CBD (<0.07%) strain, Skywalker (Aphria Inc., Lemington, ON, lot #6216), and a high-CBD (12.98%) and low-THC (0.67%) strain, Treasure Island (Aphria Inc., Lemington, ON, lot #6812), were used for *Cannabis* smoke exposure as previously established (Barnard et al., 2022; Roebuck et al., 2022). All *Cannabis* was stored in light-protected containers at room temperature. On the day of testing, whole *Cannabis* buds were ground in a standard coffee grinder for 5 sec. Then, 300 mg of the ground bud was measured and loaded into a ceramic coil. Using a 4-chamber inhalation system from La Jolla Alcohol Research, Inc., the *Cannabis* was combusted. A session started with a 5-min acclimation period, then a 1- min combustion occurred through three 5 sec ignitions with a 15 sec delay in-between each ignition. The temperature was set to 149°C, with a wattage of 60.1 W on the SV250 mod box. The smoke was then drawn into the clear Plexiglas chambers at a flow rate of 10-12 L/min. Following the 1-min combustion cycle, pumps were turned off for 1 min before they were turned back on for 13-min to gradually evacuate the smoke. Thus, the total exposure time was 15 min following initial ignition of the *Cannabis.* Rats were then moved to the testing apparatus to start the behavioral tests 20 min after the start of the combustion cycle.

### Behavioral Analysis

For validation of spontaneous WM tasks, behavioral videos were collected from an overhead perspective in black and white at a frame rate of 30 frames per second (fps) with a resolution of 720 pixels x 480 pixels (Panasonic WV-BP334 1/3” B&W). Collected videos were manually scored using a conventional stopwatch method, where the duration of interaction at each stimulus was recorded. To allow for automated behavioral analysis, behavioral videos for the *Cannabis* exposure experiment were recorded from an overhead perspective in full color at a frame rate of 30 fps and a resolution of 1080 pixels x 1080 pixels (Logitech Brio 505, Logitech). After filming, DeepLabCut (DLC) was utilized to continuously track the spatial location of eight user defined points-of-interest (Fig 3B) (Mathis et al., 2018). To train the DLC model, we randomly extracted 300 frames from 60 representative behavioral videos, with an equal representation of the IOT/DOT variation and object/odor stimuli. Manually annotated frames were used to train a deep neural network-based model to predict the spatial location of points of interest for each frame across new videos.

**Figure 3.**
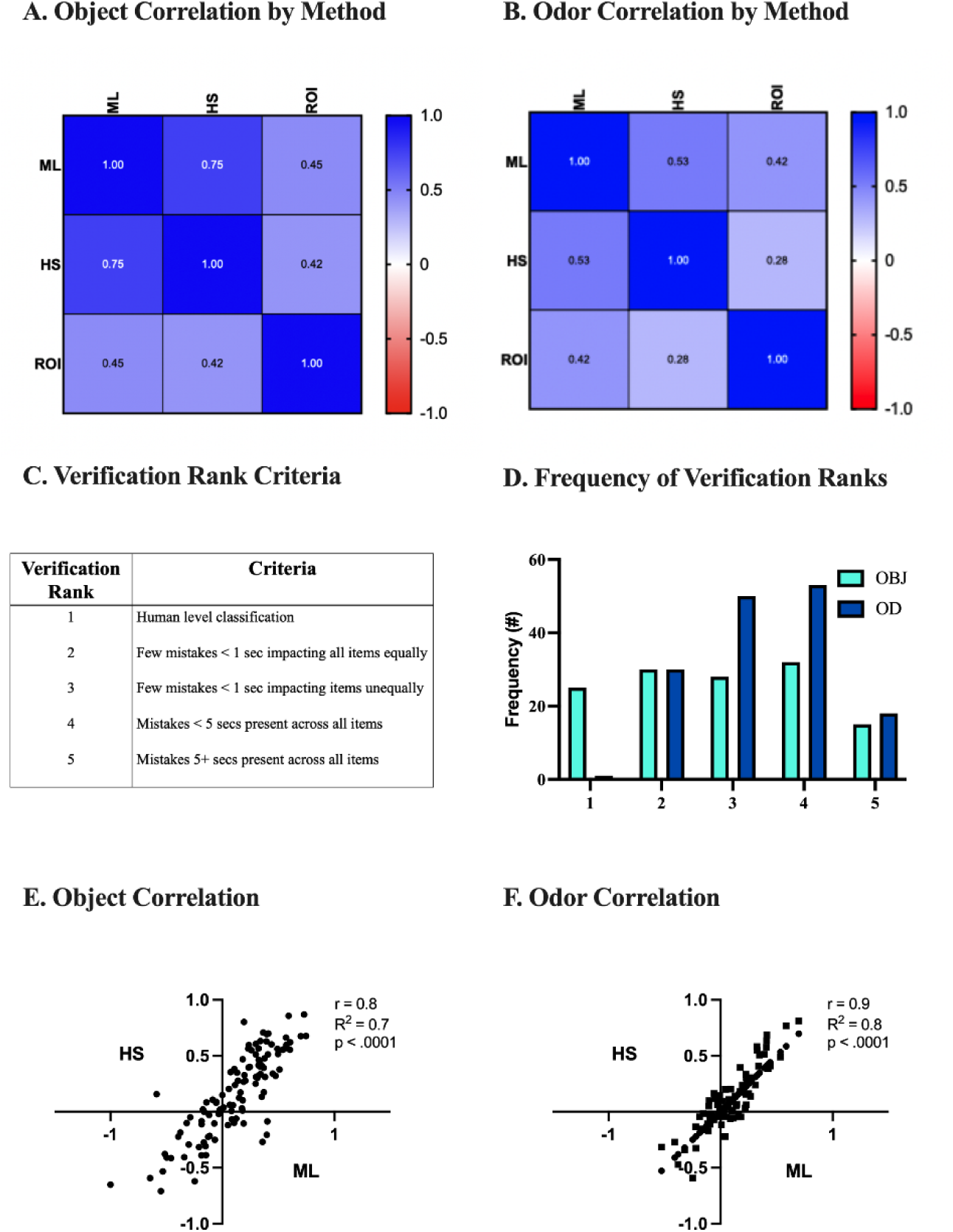
Comparison between human stopwatch and machine-learning generated output. **A** Correlation matrix between methods of quantifying rat-object interaction. Interaction times by object was quantified using each scoring method, then the correlation between interaction DR’s was assessed. **B** Correlation matrix between methods of quantifying rat-odor interaction. Interaction times by odor was quantified using each scoring method, then the correlation between interaction DR’s was assessed. **C** Criteria used to rank automated classification. Each video was manually viewed for accurate classification, where a verification rank was assigned based on objective criteria. **D** Frequency of verification rank assignment by type of stimuli. Videos with a verification rank less than three were excluded from final analysis and replaced by human stopwatch scoring. Approximately 80% of object videos and 60% of odor videos met inclusion criteria, respectively. **E** Correlation between human stopwatch and ML-generated DR’s on object videos meeting inclusion criteria, indicating a moderate-to-high correlation (r(109) = .83, p < .0001). **F** Correlation between human stopwatch and ML-generated DR’s on odor videos meeting inclusion criteria, indicating a moderate-to-high correlation (r(77) = .87, p < .0001).

We then trained a SML-based behavioral classifier to predict interaction events based on movement features extracted from pose-estimation data (Goodwin et al., 2022; Nilsson et al., 2020). Two classifiers were trained, one for each type of stimuli (Object and Odor), using 28,586 and 32,872 manually annotated frames of target “interaction” behavior, respectively (Fig 3D, Fig 3E). For the object classifier, we defined “interaction” as frames where the rat’s nose was within 2 cm of the object, while looking at and/or chewing the stimuli for a duration greater than 50 msec. For the odor classifier, “interaction” was defined as frames where the rat’s nose was within 2 cm of the top of the odor jar, while looking at and/or chewing the stimuli for a duration greater than 50 msec. To account for instances of sub-optimal SML predictions, we created a five-tiered verification rank system, where SML-generated predictions on videos with a rank less than three were replaced by human stopwatch scoring for the final analysis (Fig 3C).

### Statistical Analysis

For all analyses, the entire 5 min of the sample or test phase was analyzed. Total stimuli exploration times were calculated by taking the sum of the time spent interacting with each stimulus, as measured in sec. A discrimination ratio (DR) was calculated for each test session, which reflects the time spent with the novel stimulus compared to the average time spent with the familiar stimuli. This metric is calculated by the equation DR = (T (novel) – T (avg. familiars) / T (total)), and produces a ratio between -1 and +1, that indicates a familiar and novelty preference, respectively. A DR was also calculated for interaction bout count, while untransformed values were used to assess distance travelled and novel approach latency. Rats were excluded from the final analysis if all stimuli in the box were not visited in the sample phase, if an item was knocked over or moved, or if the video was blurry. From the test establishment experiments, 2 rats were removed from the 3-object IOT, 1 from the 3-odor IOT, 1 from the 3-odor DOT, and 1 from the 6-odor IOT. Due to missing video footage, 8 values are missing from each 3- and 6- object IOT and DOT sample phase mean ± SEM calculations. From the acute *Cannabis* exposure interaction bout duration data, 6 videos were excluded from the 6- object IOT, 2 from the 6-object DOT, 1 from the 6-odor IOT, and 2 from 6-odor DOT. From the bout count data, 7 were excluded from the 6-object IOT, 3 from the 6-object DOT, and 2 from 6- odor DOT.

Data were analyzed using GraphPad Prism 8.0.1 software. To evaluate the DR’s generated from interaction times in the test validation and establishment experiment, one-sample t-tests were used against chance (i.e., 0). To evaluate the total exploration times in the test validation and establishment experiment, two-way ANOVAs (followed by Bonferroni’s multiple comparisons test) with factors of Phase (sample vs test) and Item Count (3- vs 6-) were used. To evaluate the total exploration times following *Cannabis* smoke exposure, two-way ANOVAs (followed by Bonferroni’s multiple comparisons test) with factors of Phase (sample vs test) and Treatment (Air Control vs high-THC [Skywalker] vs high-CBD [Treasure Island]) were used. Following *Cannabis* exposure, to evaluate the DR’s and untransformed values measuring interaction time, bout count, distance travelled, and novel approach latency, one-way ANOVAs (followed by Turkey’s multiple comparisons test) with a factor of Treatment (Air Control vs high-THC vs high-CBD) were used. Lastly, to evaluate the interaction time DRs (novelty preference) against chance, one-sample t-tests against 0 were used. P values that were < or = to 0.05 were considered significant.

## Results

### Rats perform both object-and odor-based IOT and DOT tests with 3- or 6- stimuli

The 3- and 6-object IOT and DOT tests were validated for rats by adopting protocols like those used with mice (Olivito et al., 2016, 2019; Sannino et al., 2012) (Fig 1A). Rats spent significantly more time with the novel object in comparison to the familiar objects in the 3-object IOT [t(14)L= -6.29, pL< 0.001], and in the 6-object IOT [t(14)L= -5.02, pL< 0.001] test variations (Fig 1A). Rats also displayed novelty preference in the 3-object DOT [t(16)L= -5.09, pL< 0.001], and in the 6-object DOT [t(14)L= -3.94, pL< 0.001] test variations (Fig 1A). A comparison of the IOT and DOT DRs showed no differences between the 3-object [t(30) = 0.98, p = 0.36] or 6-object [t(28) = 1.40, p = 0.17] test variations (Fig 1A). All treatment groups performed better than chance (t(15) = 7.35, p < 0.0001 (3-object IOT); t(14) = 8.41, p < 0.0001 (6-object IOT); t(15) = 8.52, p < 0.0001 (3-object DOT); t(14) = 7.31, p < 0.0001 (6-object DOT) (Fig 1A).

A significant effect of Phase was seen on the total stimuli interaction time in the object IOT [F(1, 39) = 9.63, p = 0.004], with no effect of Item Count [F(1, 39) = 1.62, p = 0.21] or an interaction [F(1, 39) = 0.11, p = 0.74] present (Table 1). Rats spent more time exploring stimuli in the sample phase of the 6-object IOT than the test phase (p = 0.0054). There was also a significant effect of Phase on the total stimuli interaction time in the object DOT [F(1, 39) = 13.89, p = 0.0006], with no effect of Item Count [F(1, 39) = 3.78, p = 0.059] or an interaction [F(1, 39) = 2.61, p = 0.11] present (Table 1). In the 6-object DOT, rats spent more time exploring stimuli in the sample phase than the test phase (p = 0.0089).

**Table 1.**
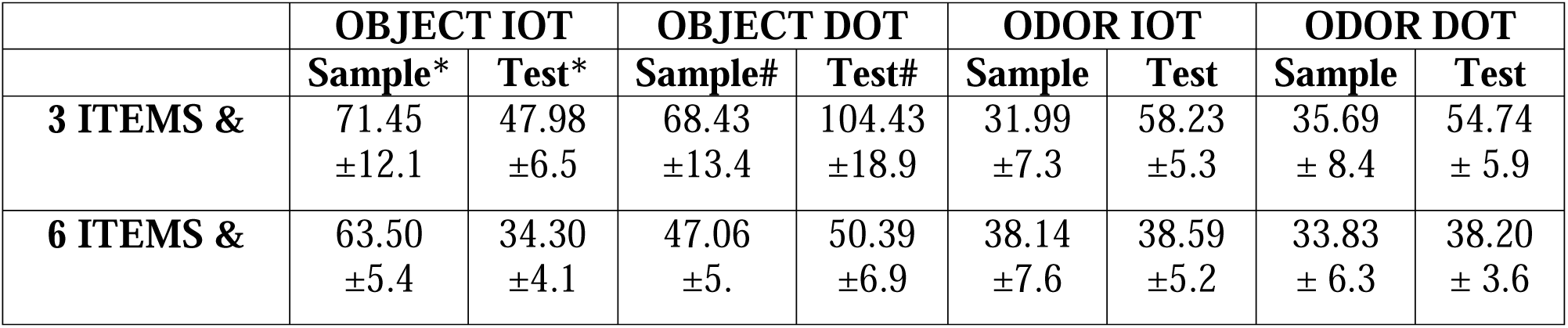
Summary of all interaction times for validation of the tests summarized in Fig 1. The mean (± SEM) for the total interaction time seen with stimuli is recorded for each sample and test phase in the different object and odor IOT and DOT variations. * Significant effect of Phase (p = .0004) on object IOT. # Significant effect of Phase (p = 0.0006) on object DOT. & Significant effect of Item Count (p = .047) on odor IOT.

In the tests with odors, rats also showed novelty preferences in the 3- and 6- odor IOT and DOT (Fig 1B). Rats spent significantly more time with the novel odor compared to the familiar odors in the 3-odor IOT [t(7)L= -1.87, pL< 0.05] and 6-odor IOT [t(10)L= -6.59, pL< 0.001] test variations (Fig 1B). Novelty preference was also demonstrated in the 3-odor DOT [t(6)L= -7.94, pL< 0.001], and in the 6-odor DOT [t(11)L= -3.92, pL< 0.01] test variations (Fig 1B). Lastly, no differences between the IOT and DOT DR’s were found in the 3-odor [t(13) = - 1.44, p = 0.17] or 6-odor [t(21) = 1.60, p = 0.12] test variations (Fig 1B). All treatment groups performed better than chance (t(7) = 5.04, p = 0.0015 (3-odor IOT); t(11) = 7.36, p < 0.0001 (6- odor IOT); t(7) = 5.40, p = 0.0010 (3-odor DOT); t(11) = 10.61, p < 0.0001 (6-odor DOT) (Fig 1B).

In the odor IOT, there was no effect of Phase on the total stimuli interaction time [F(1, 36) = 1.16, p = 0.29], but a main effect of Item Count [F(1, 36) = 4.55, p = 0.040] and a significant interaction was present [F(1, 36) = 4.24, p = 0.047] (Table 1). Rats spent more time exploring odors in the sample phase of the 6-odor IOT than in the 3-odor IOT (p = 0.031). In the odor DOT, there was no main effect of Phase [F(1, 36) = 2.34, p = 0.14], Item Count [F(1, 36) = 3.79, p = 0.06] or an interaction [F (1, 36) = 1.49, p = 0.23] present (Table 1).

### Combining automated and human stopwatch scoring is a valid behavioral quantification approach

To quantify rat behavior following *Cannabis* smoke exposure using the HYB scoring method, we created a video set of 288 test phase videos of the 6-stimuli task variations. Sample phase videos were all manually scored, where inclusion criterion was applied as described above, and included test phase videos were analyzed using our automated behavioral quantification pipeline.

To assess the accuracy of model predictions for both pose-estimation and behavioral classification, we utilized software native performance metrics that compare machine-generated predictions to manual annotation. The spatial coordinates of human annotated and machine-predicted points-of-interest differed by a mean Euclidian distance of 4.89 pixels on videos within the model training set and 4.35 p on test videos. Pose-estimation quality was further assessed by calculating the average prediction confidence for each point-of-interest by video (Supplementary Figure 2). We found that the average prediction confidence ranged between 92.8% and 97.4% by point-of-interest, where no significant differences were observed between object-based and odor-based videos. Behavioral classifier performance was evaluated by a series of confusion matrices (Fig 3D, Fig 3E) that report the precision, recall, and combined F1 score for each model. In short, both classifiers demonstrate high precision and recall (object F1 = 0.927, odor F1 = 0.897) when assessed by comparing manual annotation to classifier predictions on randomly selected test video frames. However, when classifier performance was assessed by comparing predictions on randomly selected interaction bouts, object classifier performance changed marginally (F1 = 0.93), but odor classifier performance decreased markedly (F1 = 0.63). For both the object and odor classifiers, the behavioral features most heavily weighted for model predictions include distance to stimuli, nose movements, region-of-interest, and spatial dynamics between points-of-interest (Fig 3C). Additional detail regarding model training and assessments can be found in supplementary materials.

To verify the reliability of SML-generated predictions relative to traditional stopwatch-based and automated ROI-based scoring, we conducted a three-way correlational analysis on generated interaction DR’s (Fig 3A, B). We found that, across stimuli, SML-generated predictions were more highly correlated with human stopwatch scoring than ROI-based scoring; however, SML-generated predictions were more highly correlated with human stopwatch scoring for object interaction (r = 0.75) relative to odor interaction (r = 0.53). Additionally, we found that, across stimuli, ROI-based scoring held a weaker correlation relative to both human stopwatch scoring (object: r = 0.42, odor: r = 0.28) and SML-generated (object: r = 0.45, odor: r = 0.42) interaction DR’s. To account for instances where SML predictions significantly differ from human stopwatch scoring, we created a five-tiered verification rank system, where SML-generated predictions on videos with a rank less than three were replaced by human stopwatch scoring for the final analysis. Upon visual inspection of SML-generated predictions, we found that 80% of object-based videos met inclusion criteria, while only 60% of odor-based videos met inclusion criteria. To justify supplementing human stopwatch scoring for sub-optimal SML-generated predictions, we conducted a correlational analysis between human stopwatch scoring and SML interaction DR’s only on videos which met inclusion criteria. We found that human stopwatch scoring and SML interaction DR’s were moderately-to-highly correlated (Fig 3E: r = 0.83, Fig 3F: r = 0.87) across stimuli type.

### High-THC, but not high-CBD, Cannabis smoke exposure impairs novelty preference for high-(DOT) cognitive loads using object stimuli

The impact of *Cannabis* smoke exposure on locomotion was evaluated first. We found no main effects of Treatment on distance in either the 6-object IOT [F(2, 70) = 0.58, p = 0.56], or in the 6-object DOT [F(2, 67) = 0.30, p = 0.74] tests (Fig 4C). Next, we investigated novel approach latency values, defined as the interval between rats being placed into the experimental arena and interacting with the novel object. No effect of Treatment on novel approach latency values was observed in either the 6-object IOT [F(2, 70) = 0.77, p = 0.46] or the 6-object DOT [F(2, 67) = 0.076, p = 0.93] tests (Fig 4D). Then, to examine if the rat visited the novel object at a higher frequency than familiar objects, we evaluated the interaction bout DR’s (Fig 4E). Here, we showed a significant main effect of Treatment in the 6-object IOT [F(2, 64) = 8.05, p < 0.001] test, as the Air Control (p = 0.001) and high-THC (p = 0.01) groups were different from the high-CBD group. However, we failed to find a main effect of Treatment on bout count DR’s in the 6-object DOT [F(2,64) = 0.96, p= 0.39] test (Fig 4E). Then, interaction bout duration DR’s were investigated to examine if novelty preference was impacted by treatment within each test variation. No effect of Treatment in the 6-object IOT [F(2, 61) = 0.85, p = 0.43] test was found (Fig 4F). A main effect of Treatment was present in the 6-object DOT [F(2, 63) = 3.75, p = 0.03] test, with a significant difference seen between the Air Control and high-THC groups after a Tukey’s multiple comparisons test (p = 0.04) (Fig 4F). Most treatment groups performed better than chance (t(23) = 3.15, p = 0.004 (IOT-Air Control); t(19) = 2.24, p = 0.037 (IOT-high-THC); t(19) = 4.27, p = 0.0004 (IOT-high-CBD); t(18) = 3.29, p = 0.004 (DOT-Air Control); t(24) = 2.14, p = 0.042 (DOT-high-CBD)) except for the high-THC group in the 6-object DOT test (t(22) = 0.66, p = 0.51) (Fig 4F).

**Figure 4.**
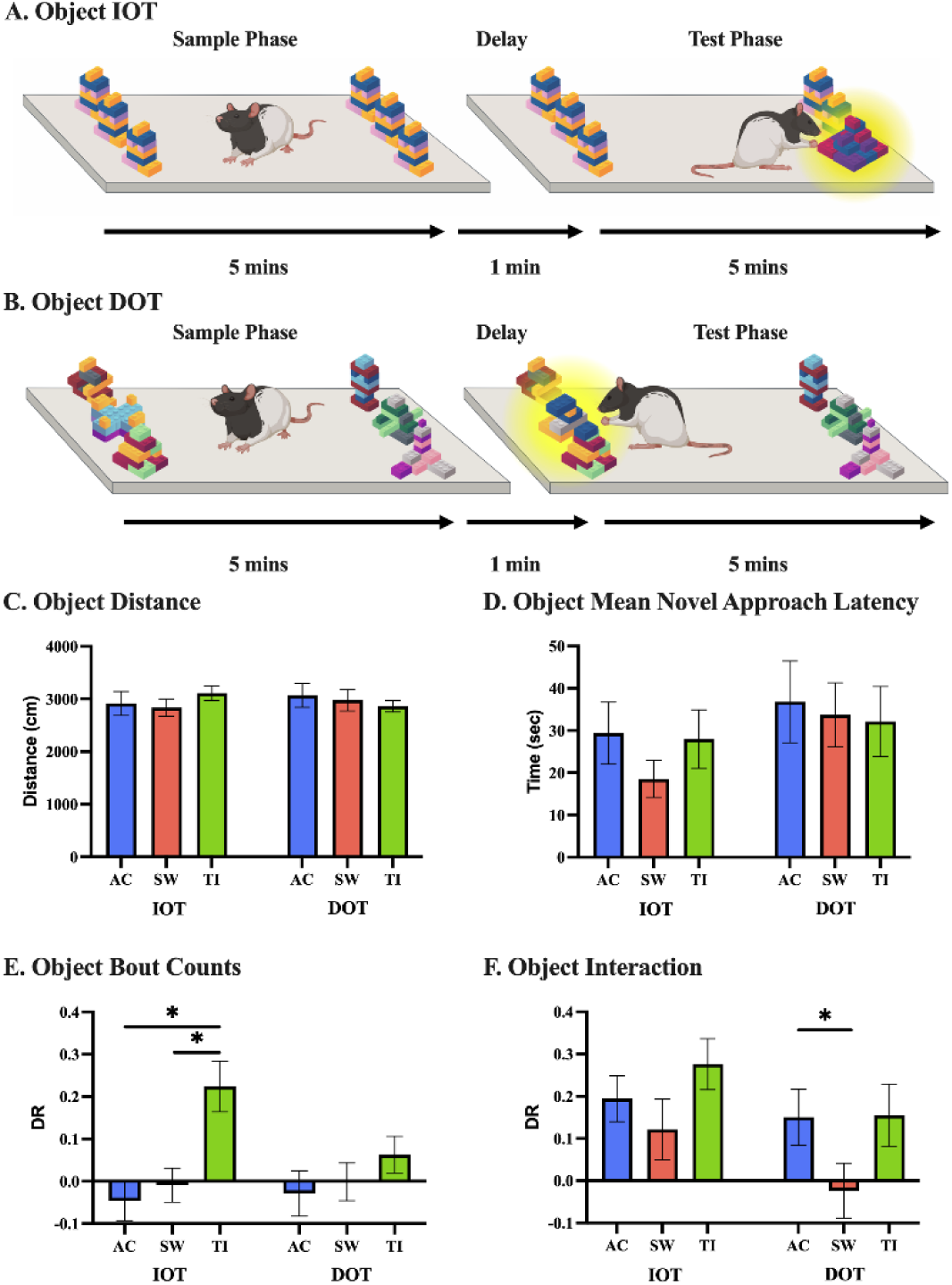
High-THC *Cannabis* smoke exposure impacts novelty preference under high-(DOT) cognitive loads using object stimuli, with no impact on distance travelled, frequency of item visitation, or approach latencies. A An example object IOT is visualized, showing 6 identical objects in the sample phase, with a novel object introduced after a 1-minute delay in the test phase. B An object DOT variation is shown, with an identical task progression, but instead starts with 6 different objects in the sample phase. C The distance travelled (cm) in the 6-object IOT (n = 72) and 6-object DOT (n = 69) variations is comparable across treatment groups. D The mean novel approach latency in the 6-object IOT (n = 72) and 6-object DOT (n = 69) variations is shown to be consistent between treatment groups. E To illustrate the frequency of visitations to the novel object in comparison to the familiar objects, bout counts are visualized using a discrimination ratio. A preference for novel visitations is seen in the 6-object IOT (n = 65) AC and SW groups, with no difference in item visitations in the 6-object DOT (n = 66). F Interaction measured as time spent with an object was generated using the human-machine hybrid scoring approach and visualized using a discrimination ratio for both variations using object stimuli. No difference in treatment groups is seen in the 6-object IOT (n = 64). In the 6-object DOT (n = 66), a significant decrease in novelty preference is seen in the SW group in contrast to the AC group (p = .04). Data represents mean ± SEM. *pL<L0.05. Abbreviations: High-THC *Cannabis* smoke (SW), high-CBD *Cannabis* smoke (TI), Air Control (AC). This figure was created using BioRender.com.

When assessing total stimuli interaction time, a main effect of Treatment [F(2,129) = 4.07, p = 0.019], and of Phase [F(1, 129) = 6.45, p = 0.012] was seen in the 6-object IOT, with no significant interaction [F(2, 129) = 0.49, p = 0.62] (Table 2). In the 6-object DOT, there was a main effect of Phase on total stimuli interaction time [F(1, 135) = 7.87, p = 0.0058], with no main effect of Treatment [F(2, 135) = 1.81, p = 0.17] or an interaction [F(2, 135) = 0.75, p = 0.47] (Table 2).

**Table 2.** Summary of all interaction times for tests with *Cannabis* summarized in Fig 2-5. The mean (± SEM) for the total interaction time seen with stimuli is recorded for the sample and test phases in the different 6- object and 6- odor IOT and DOT variations across the Air Control, high-THC, and high-CBD treatment groups. * Significant effect of Treatment (p = 0.019) and of Phase (p = 0.012) on object IOT. # Significant effect of Phase (p = 0.0058) on object DOT. & Significant effect of Treatment (p = 0.025) and Phase (p = 0.0004) on odor IOT. % Significant effect of Phase (p = 0.0019) on odor DOT.

|  | OBJECT IOT |  | OBJECT DOT |  | ODOR IOT |  | ODOR DOT |  |
| --- | --- | --- | --- | --- | --- | --- | --- | --- |
|  | Sample* | Test* | Sample# | Test# | Sample& | Test& | Sample% | Test% |
| **Air Control**<br>**\*&** | 36.21<br>±2.9 | 42.93<br>±4.0 | 35.61<br>±3.2 | 39.23<br>±3.4 | 37.75<br>±2.8 | 47.78<br>±5.8 | 39.16<br>±3.1 | 50.12<br>±5.6 |
| **high-THC**<br>**\*&** | 36.01<br>±3.7 | 46.90<br>±4.1 | 39.65<br>±3.5 | 49.72<br>±4.6 | 34.27<br>±3.1 | 57.94<br>±4.8 | 35.29<br>±2.8 | 55.27<br>±6.5 |
| **high-CBD**<br>**\*&** | 30.09<br>±3.0 | 33.97<br>±2.7 | 33.9<br>±3.1 | 46.96<br>±4.2 | 31.54<br>±2 | 36.93<br>±5.5 | 40.54<br>±3.4 | 48.03<br>±6.1 |

### High-THC, but not high-CBD, Cannabis smoke exposure impairs novelty preference for high-(DOT) and low-(IOT) cognitive loads using odor stimuli

*Cannabis* smoke exposure did not impact the distance travelled by the rat in either the 6- odor IOT [F(2, 77) = 0.36, p = 0.70], or in the 6-odor DOT [F(2, 71) = 0.87, p = 0.42] tests (Fig 5C). As well, no effect of Treatment in the 6-odor IOT [F(2, 77) = 0.036, p = 0.70], or in the 6- odor DOT [F(2, 71) = 0.87, p = 0.42] tests were seen when investigating novel approach latency (Fig 5D). Interaction bout DR’s were also determined to be unaffected by *Cannabis* exposure with no effect of Treatment in the 6-odor IOT [F(2, 77) = 1.46, p = 0.24], and the 6-odor DOT [F (2, 70) = 2. 19, p= 0.12] tests (Fig 5E). When examining interaction bout duration DRs, an effect of Treatment in the 6-odor IOT [F(2, 73) = 3.54, p = 0.034] test was seen, with a significant difference present between the Air Control and high-THC groups (Tukey’s multiple comparisons test, p = 0.046) (Fig 5F). A main effect of Treatment was also present in the 6-odor DOT [F(2, 71) = 4.3, p = 0.017] test, with a significant difference seen between the Air Control and high-THC groups (p = 0.024) and between high-THC and high-CBD groups (p = 0.046) after a Tukey’s multiple comparisons test (Fig 5F). Most treatment groups performed better than chance (t(25) = 5.90, p < 0.001 (IOT-Air Control); t(22) = 2.47, p = 0.022 (IOT-high-CBD); t(23) = 3.45, p = 0.002 (DOT-Air Control); t(27) = 2.25, p = 0.033 (DOT-high-CBD)) except for the high-THC group in the 6-odor IOT (t(26) = 0.47, p = 0.64) and 6-odor DOT tests (t(21) = 1.0, p = 0.33) (Fig 5F).

**Figure 5.**
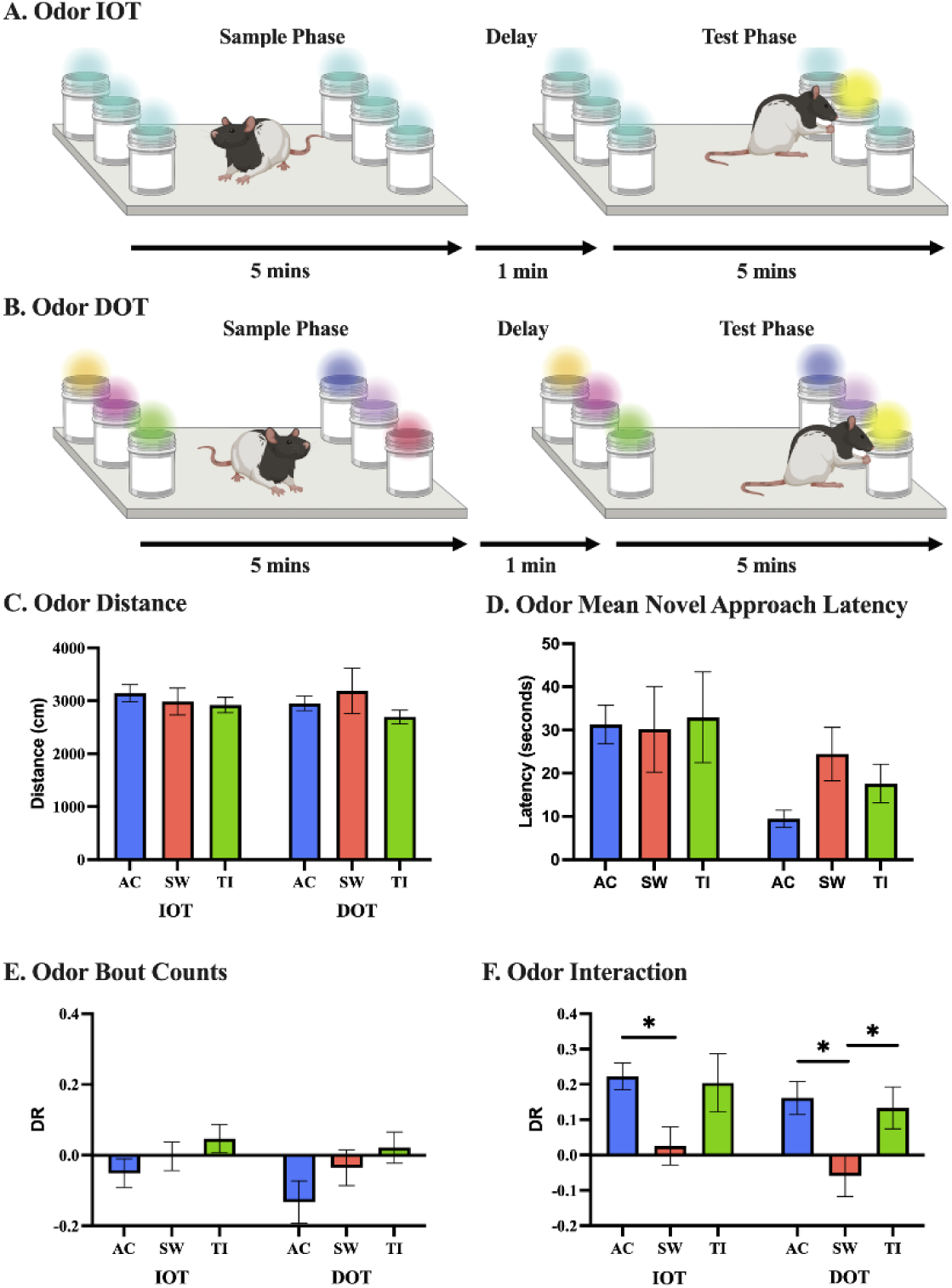
High-THC *Cannabis* smoke exposure impacts novelty preference under high-(DOT) and low-(IOT) cognitive loads using odor stimuli, with no impact on distance travelled, frequency of item visitation, or approach latencies. A An example odor IOT variation is visualized, showing 6 identical items in the sample phase, with a novel odor introduced after a 1-minute delay in the test phase. B An odor DOT variation is shown, with an identical task progression, but instead starts with 6 different odors in the sample phase. C Distance travelled (cm) in the 6-odor IOT (n = 79) and 6-odor DOT (n = 73) variations is comparable across treatment groups. D The mean novel approach latency in the 6-odor IOT (n = 79) and 6-odor DOT (n = 73) variations is shown to be consistent between treatment groups. E To illustrate the frequency of visitations to the novel odor in comparison to the familiar odors, bout counts are visualized using a discrimination ratio. No differences between treatment groups or 6-odor IOT (n = 79) and 6-odor DOT (n = 73) is seen. F Interaction measured as time spent with an odor was generated using the human-machine hybrid scoring approach and visualized using a discrimination ratio for both variations using odor stimuli. In the 6-odor IOT (n = 75), a significant decrease in novelty preference is seen in the AC group in comparison to the SW group (p = .046). Whereas in the 6-odor DOT (n = 73), a significant decrease in novelty preference is seen in the SW group from both the AC (p = .023) and TI (p = .046) groups. Data represents mean ± SEM. *pL<L0.05. Abbreviations: High-THC *Cannabis* smoke (SW), high-CBD *Cannabis* smoke (TI), Air Control (AC). This figure was created using BioRender.com.

In the 6-odor IOT, a main effect of Treatment [F(2,142) = 3.78, p = 0.025], and of Phase [F(1, 142) = 12.90, p = 0.0004] was seen, with no significant interaction [F(2, 142) = 2.27, p = 0.11] (Table 2). Rats spent more time exploring stimuli in the Air Control sample phase than in the high-THC test phase (p = 0.017). As well, rats explored stimuli more in the sample phase than in the test phase following high-THC (p = 0.0035), while spending more time exploring stimuli in the test phase following high-THC smoke exposure than following high-CBD smoke exposure (p = 0.009). In the 6-odor DOT, there was a main effect of Phase on total stimuli interaction time [F(1, 134) = 10.01, p = 0.0019], with no main effect of Treatment [F(2, 134) = 0.021, p = 0.98] or an interaction [F(2, 134) = 0.85, p = 0.43] (Table 2).

## Discussion

In the present study, we showed that rats display novelty preference in both the IOT and DOT with 3 and 6 objects, similar to previous findings using objects in mice (Olivito et al., 2016, 2019; Sannino et al., 2012). We also demonstrate, for the first time, that rats exhibit novelty preference with 3 and 6 odor stimuli, as measured in the IOT and DOT tests (Fig 1B). Overall, rats spent more time exploring stimuli in the sample phases of the 6 item IOT and DOT tests compared to the test phases, with stimuli-specific differences (Table 1). Following high-THC *Cannabis* smoke exposure in the tests with objects, a significant decrease in novelty preference was seen in the 6-object DOT, but not in the 6-object IOT (Fig 4F). However, for odor-based tests, we observed novelty preference impairments for high-and low-cognitive loads (Fig 5F). No notable treatment effect on total stimuli exploration time was present in the 6-object IOT tests, but a significant increase in stimuli exploration time was seen in the test phase of the 6- object DOT for all treatments (Table 2). In the 6- odor IOT, rats explored stimuli less in the sample phase compared to the test phase following high-THC *Cannabis* smoke exposure, with no notable effects in the 6-odor DOT (Table 2). Taken together, these findings suggest that *Cannabis* smoke exposure impacts novelty preference in a load-dependent and stimuli-specific manner.

### Rats demonstrate novelty preference in both the object-and odor-based IOT and DOT tests

In the validation and establishment experiments, rats demonstrated pronounced novelty preference in all test variations (Fig 1). The preferential interaction with novel stimuli compared to familiar stimuli after a brief delay suggests that WM processes are involved in both object and odor-based tests (Sannino et al., 2012; Shrager et al., 2008; van Vugt et al., 2017). The varying cognitive loads between the IOT and DOT test variations also present the opportunity to examine WMC (Sannino et al., 2012; Shrager et al., 2008). Rats explored the object stimuli a comparable amount between test variations and with varying numbers of stimuli (Table 1). Rats did, however, spend significantly less time exploring objects in the test phase of the 6-object DOT compared to the sample phase (Table 1). As the test phase progressed, rats would have had increasing familiarization with all items in the test phase, which may explain the decreased total exploration times (Broadbent et al., 2010). Interestingly, there were no notable differences in the total stimuli interaction times between the 3-odor and 6-odor variations, indicating that while the total time rats spent exploring stimuli was the same, the time spent exploring each individual stimulus in the 6-item variation was about half of that for the 3-item variation (Table 1). In future experiments, it would be interesting to assess novelty preferences and exploration preferences in test with more than 6 stimuli, as has been reported for objects in mice (Sannino et al., 2012).

The IOT and DOT tests present a unique method to study novelty preference in a spontaneous, simple, and cost-effective manner. The tests do not require rodents to apply learned rules or procedures, eliminating extensive training and needing minimal researcher involvement. The tests also evoke minimal stress in rodents, as spontaneous exploration tests do not have the prerequisite of typical food-restriction protocols to induce reward-driven performance. Performance on the object tests likely engage a combination of visual and tactile recognition memory, but as the object stimuli were constructed with LEGO™ blocks of similar size, identical smooth textures, and sharp corners, the tests were likely biased to engage visual recognition memory. The object-based test may engage visual, perirhinal, medial prefrontal, parietal, and entorhinal cortices, as well as the hippocampus and thalamus to enable the object-based recognition memory across a delay (Barker et al., 2007; Cazakoff & Howland, 2011; Churchwell & Kesner, 2011; Creighton et al., 2018; Dere et al., 2007; Fernandez & Tendolkar, 2006; Hannesson et al., 2004; Peters et al., 2013; Sugita et al., 2015; Winters et al., 2004). The odor stimuli primarily engage odor-based recognition as identical opaque glass jars were used in the tests. A circuit including piriform, entorhinal, medial prefrontal, and orbitofrontal cortices, along with hippocampus may be involved in the odor-based memory across a delay (Alvarez & Eichenbaum, 2002; Davies et al., 2013; Mouly & Sullivan, 2010; Ramus & Eichenbaum, 2000; Sandini et al., 2020).

To examine the brain regions and neural mechanisms underlying WM and WMC in different contexts, a variety of behavioral tasks can be employed. To examine visuospatial WM and WMC across a time delay, tasks like the Radial-Arm Maze (RAM), Barnes Maze, and operant delayed nonmatching-to-sample (DNMTS) and delayed-match (DMTS) tasks have been used (Barnard et al., 2022; Cowan, 2010; Daneman & Carpenter, 1980; Dudchenko, 2004; Dudchenko et al., 2013; Kirchner, 1958; Oomen et al., 2013; Scott et al., 2020; Vorhees & Williams, 2014; Wilhelm et al., 2013). To study odor based WMC, the Odor Span Task (OST) and tests that employ a nonmatch-to-sample-rules have often successfully been used (Dudchenko et al., 2000; Scott et al., 2020). Although tasks like RAM, OST, and DNMTS tasks measure WMC, these tasks require food restriction, extensive training, and heavy researcher involvement. Spontaneous recognition tests circumvent these weaknesses by measuring novelty preference that can be used to infer WM and WMC. These spontaneous IOT and DOT tests also boast the flexibility to study novelty preference in a variety of different species like mice (Bevins & Besheer, 2006; Lueptow, 2017). Validating the odor-based spontaneous tests in mice would be a worthwhile future experiment considering mice can be highly cost-effective, and there is an abundance of genetic resources to model various disease-and disorder-like states that could provide pre-clinical insight on WM and WMC deficits in humans.

### High-THC, but not high-CBD, Cannabis smoke exposure impairs novelty preferences for both object and odor stimuli

To evaluate the effects of *Cannabis* smoke exposure on WMC, we used the HYB scoring approach to assess novelty preference in the object and odor based IOT and DOT tests. Novelty preference was primarily indicated by interaction bout duration, as it was not predicted by interaction bout count and novel approach latency. Following high-THC *Cannabis* smoke exposure in the tests with objects, a significant decrease in novelty preference was seen in the 6- object DOT, but not in the 6-object IOT (Fig 4F). For odor-based tests, an impairment in novelty preference was observed in both the IOT and DOT following high-THC *Cannabis* smoke exposure (Fig 5F). In all tests, novelty preference was similar between the Air Control and high-CBD *Cannabis* smoke groups. Additionally, no differences in locomotion were observed between all treatment groups. The increased total stimuli exploration time in the sample phases of the object DOT compared to the test phases likely indicates familiarity with the items in the test phase that were previously presented during the sample phase (Broadbent et al., 2010). Interestingly, in the 6- odor IOT, there was lower stimuli exploration time in the sample phase compared to the test phase following high-THC *Cannabis* smoke exposure (Table 2).

Overall, the deficits in novelty preference following high-THC *Cannabis* smoke exposure in both the object and odor-based tests are likely attributable to the THC potency, and not to smoke alone. As behavioral testing was initiated 20 min following *Cannabis* smoke exposure, plasma and brain THC concentration would have been near its peak in the rats (Baglot et al., 2021; Barnard et al., 2022; Hložek et al., 2017; Moore et al., 2022; Ravula et al., 2019). Analysis of rat plasma following an identical *Cannabis* smoke exposure paradigm revealed levels of 14.55 ± 1.59 ng/mL with a small amount of CBD (1.98 ± 0.38 ng/mL) 30 min after smoke exposure (Barnard et al., 2022). After high-CBD smoke exposure, negligible amounts of THC were found in plasma, along with 4.47 ± 1.15 ng/mL of CBD (Barnard et al., 2022). Thus, the current smoke exposure protocol increases blood plasma levels of THC to the lower end of what is typically observed in humans following *Cannabis* cigarette consumption (Grotenhermen, 2003; Huestis, 2007; Huestis et al., 1992; Newmeyer et al., 2016; Moore et al., 2022; Ramaekers et al., 2009). Although the THC plasma levels in rats were comparably low, we were still able to observe the impact of *Cannabis* exposure on WMC.

The different THC-induced novelty preference impairments between objects and odors may be due to the varying neural circuits underlying stimulus perception and integration (Constantinidis & Klingberg, 2016; Eriksson et al., 2015; Fernandez & Tendolkar, 2006; Galizio, 2016; Mouly & Sullivan, 2010). Under low cognitive loads (IOT), treatment does not impact object novelty preference, consistent with unperturbed WM performance previously observed in a 2-item novel object recognition (NOR) test following chronic exposure to 5.6% THC *Cannabis* cigarettes (Bruijnzeel et al., 2016). However, it is possible that the sample was underpowered and that a THC-induced deficit, as seen in the 6-odor IOT, was not detected. The novelty preference deficits observed following high-THC *Cannabis* exposure in the 6-odor IOT also might have been affected by the decreased exploration time in the sample phase. Lastly, the similar THC-induced deficits in the object and odor DOT could be due to sensitivity of the WM subconstructs evoked under high cognitive loads to *Cannabis* exposure (Barch & Smith, 2008).

### The case for, and caveats of, SML-based behavioral analysis at scale

Automated behavioral analysis represents a potential paradigm shift in the way behavioral data are generated and shared (Mathis et al., 2020). In the present study, we demonstrate the case for, and caveats of, using a SML-based analysis method for complex behavior at scale. In short, pose-estimation data was used to train two behavioral classifiers to predict interaction events with object and odors. We found that SML-generated predictions were more strongly correlated with human stopwatch than ROI-based scoring; however, we observed that SML-generated predictions were more highly correlated with human stopwatch-based scoring for object stimuli than for odor stimuli. Upon visual inspection of SML-generated predictions, a near 30% increase in the proportion of excluded SML-based odor interaction DR’s is striking given that each classifier was trained on the same amount of training frames, used identical algorithm hyperparameters, and was trained to assess a similar behavioral phenotype. We propose that this difference may be explained by divergent operational definitions of interaction in object and odor tests. Rat-object events encompassed interaction along the entire height of the object, while rat-odor interaction was only counted at a narrow space around the lid of the mason jar. As we employed a 2-dimensional (2D) pose-estimation approach, movements along the height of stimuli were not well captured, potentially leading to sub-optimal predictions and grounds for exclusion. While classifiers trained on 2D pose-estimation data show reliability on classifying behaviors restricted to single-plane spatiotemporal movements, recent studies of complex behaviors, such as self-grooming, generally train classifiers on 3D pose-estimation data to better capture the entirety of a movement and to minimize occlusion (Marshall et al., 2021, 2022; Minkowicz et al., 2023; Newton et al., 2023). To fully capture behaviors of interest, researchers utilizing automated behavioral analysis should be cognisant of the angle, and number, of camera perspectives used during filming (Luxem et al., 2022). Additionally, it is essential to include a diversity of training examples for pose-estimation and behavioral classifier models, as a high degree of diversity in a training set will lead to a high degree of generalizability. Within the present study, our DLC model did not generalize well to behavioral videos filmed for task validation, likely due to divergent resolution dimensions and color contrast that was not well represented in the model training set. Software native performance metrics for both behavioral classifiers closely mirror those reported in published studies utilizing SML-based analysis; however, manual verification of predictions revealed significant instances of misclassification (Newton et al., 2023; Winters et al., 2022). We contend that supplementing classifier performance metrics with correlational analysis and verification steps are best practices when conducting scaled automated behavioral analysis.

## Conclusion

Using the novel spontaneous tests and the HYB scoring method, the impact of acute exposure to high-THC or high-CBD *Cannabis* smoke was evaluated. We show impairment in object-based novelty preference after high-, but not high-CBD *Cannabis* smoke exposure under a high-cognitive load. As well, we show deficits in odor-based novelty preference following high-THC *Cannabis* smoke exposure under both low-and high-cognitive loads. Ultimately, these data indicate that *Cannabis* smoke exposure impacts novelty preference in a load-dependent, and stimuli-specific manner.

## Supporting information

Supplemental machine learning methods

## Acknowledgements and funding sources

Funding for these experiments was obtained from the University of Saskatchewan College of Medicine and the Natural Sciences and Engineering Research Council of Canada (NSERC) to JGH. ILB and TJO were supported by scholarships from NSERC. JCA was supported by the University of Saskatchewan College of Medicine. The authors wish to thank Morgan Schatz for initial pilot research on the behavioral tests reported in this paper, and Killian Stacey for contributing to the implementation of automated behavioural analysis methods.

