## Supplemental machine learning methods for "High-THC *Cannabis* smoke impairs working memory capacity in spontaneous tests of novelty preference for objects and odors in rats"

**GitHub Link:** [**https://github.com/HowlandLab/ILBTJO_NODB_SimBA_2023**](https://github.com/HowlandLab/ILBTJO_NODB_SimBA_2023)

**Video Recording and Pre-processing:** Behavioral videos were recorded in a cuboidal experimental apparatus, colored white to maximize contrast between the subject and the apparatus. Videos were recorded from an overhead perspective at a frame rate of 30fps and a resolution of 1080p x 1080p (Logitech Brio 505, Logitech). To further standardize behavioral videos, we used the “batch preprocessing” module within Simple Behavioral Analysis (SimBA) to crop videos to only include the apparatus, to ensure standardized resolution and frame rate, and to the trim video length to desired experimental phases. Additionally, we chose to film all videos in a .mp4 format as this format is generally compatible with open-source video analysis software.

**Pose-estimation Dataset and Training:** DeepLabCut (2.2.3) was utilized to continuously track the spatial location of eight user defined points-of-interest. Here, 300 frames were randomly extracted from 60 representative videos, where included videos were counterbalanced by phase (IOT/DOT) and modality (odor, object). Next, each frame was manually annotated, where a human annotator placed digital points-of-interest on the rat (Fig 2B, Supplementary Fig 1). A pre-trained ResNet-50 convolutional neural network (CNN) was then trained on 95% of annotated frames for 200,000 iterations, where 5% of frames were reserved for model assessment. After training, we analyzed the CNN learning curve to select an optimal model that performs well on both test and train data. Pose-estimation data was extracted from videos using a model trained for 80,000 iterations, which represents the iteration where test error is minimized, and the training error is saturated. Our model produced a training error of 4.89 and a test error of 4.35 using the default hyperparameters, without a p-cutoff filter applied. Finally, pose-estimation tracking files were filtered using the DeepLabCut native median filter model.

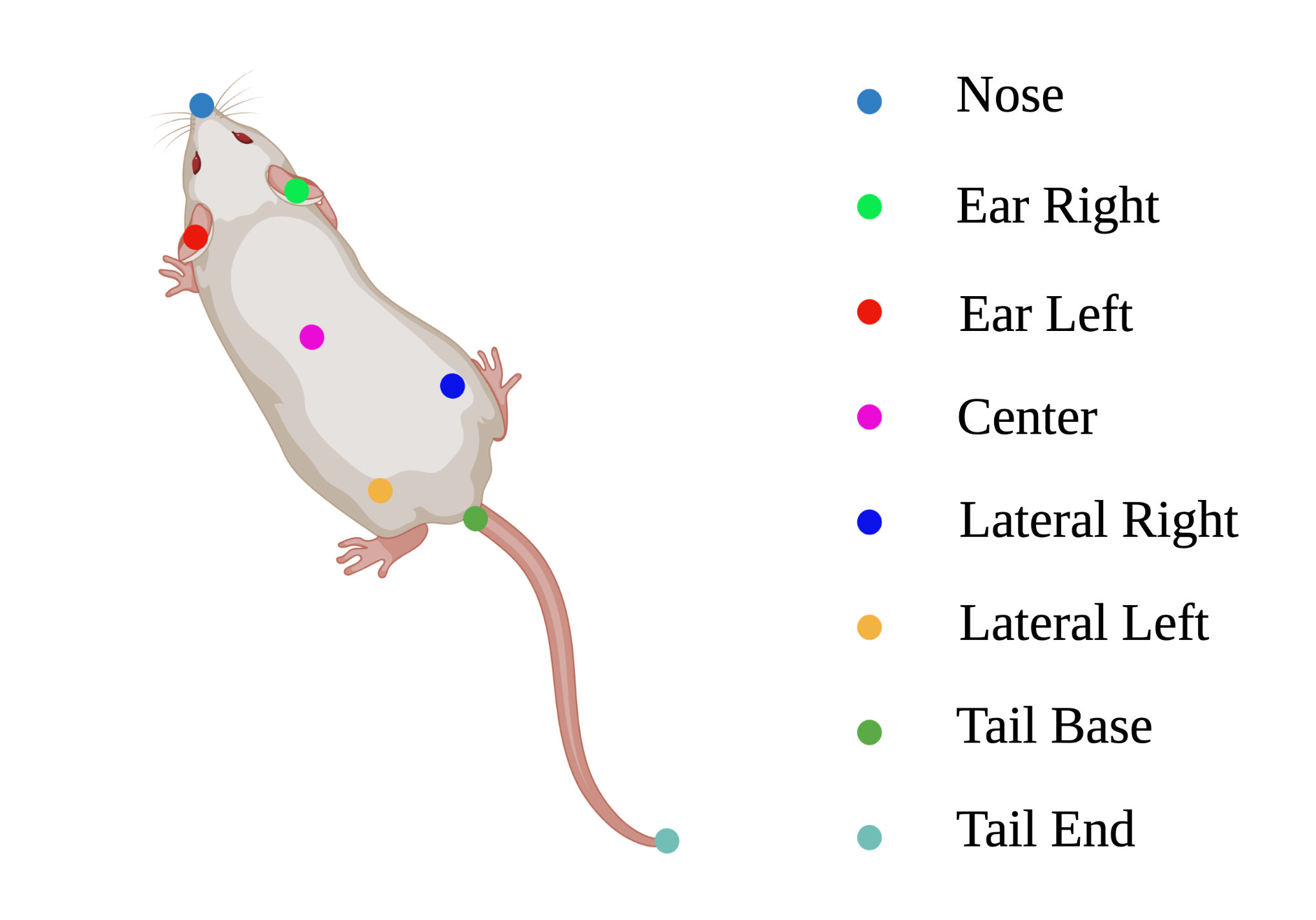
**Supplementary Fig 1**. Visualization of points-of-interest used for this experiment. We chose the number and position of points in accordance with the SimBA eight-point configuration. SimBA requires a standardized and specific position (and number) of points. Users should decide what SimBA configuration will be used (single animal, multi animal, point number) prior to network training with DeepLabCut. This figure was created using BioRender.

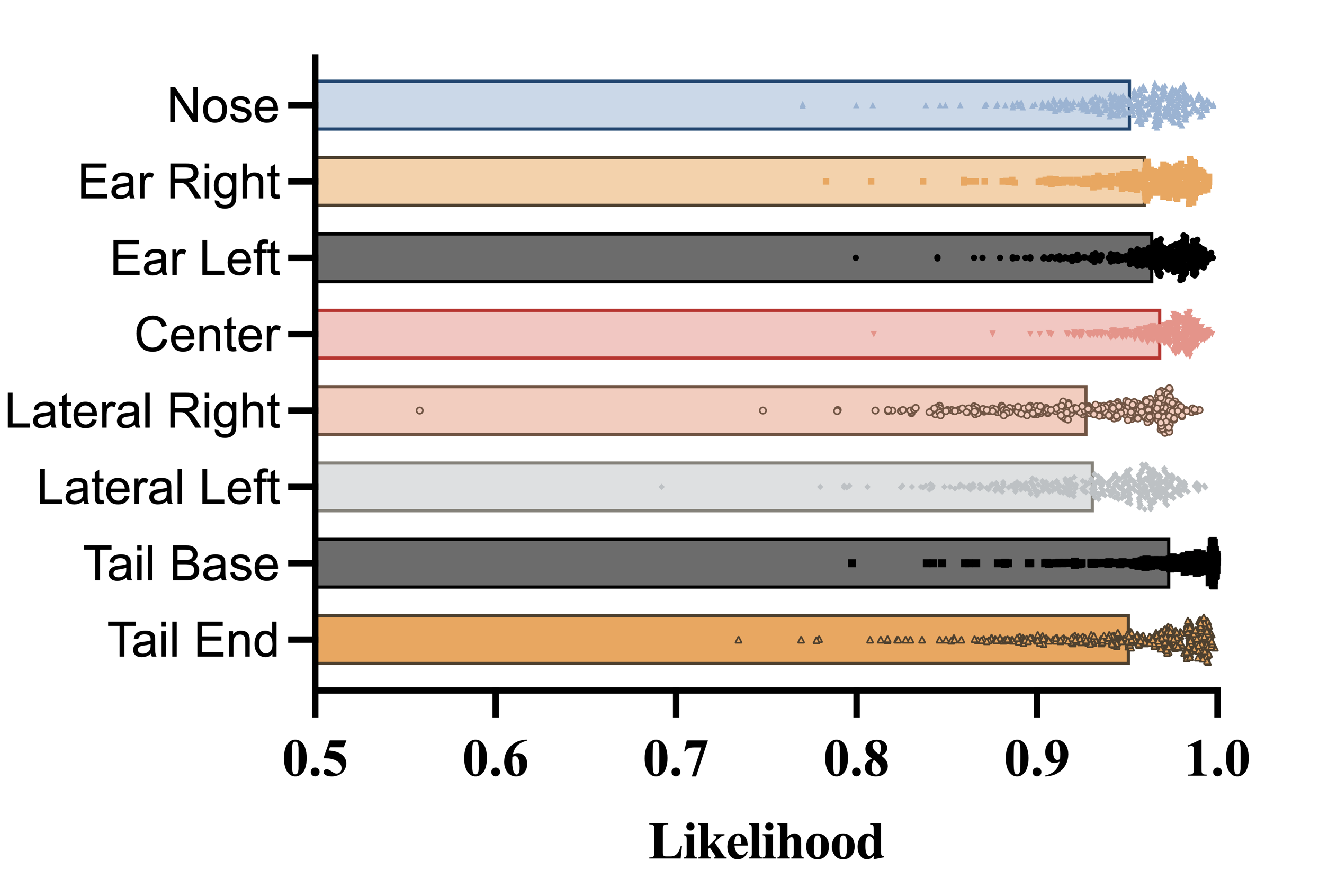

**Supplementary Fig 2.** Mean tracking confidence for each point-of-interest, by video. To calculate the mean tracking confidence for each video, the average of the likelihood column associated with each point of interest was calculated.

It is important to note that annotated training frames for this experiment were added to an existing DLC project (training set = ~1,000 annotated frames). As the CNN was pretrained to predict the spatial position of key points, and all videos were filmed within an identical experimental apparatus, the number of additional required annotated frames to acquire high-fidelity pose-estimation data for the present experiment was likely lower than if the CNN was trained from scratch. The DLC model file used for analysis is freely available on GitHub, and any additional training data will be freely supplied upon request.

**Behavioral Classifier Dataset and Training:** Classifier training was completed using the eight-point classical tracking version of the SimBA pipeline (SimBA-UW-tf-dev = 1.32.2). We trained two classifiers, one for object-based stimuli and one for odor-based stimuli, to predict interaction events across task variation. For each classifier, the training dataset consisted of user-annotated frames from ~30 five-minute videos, where each frame was assigned a binary label of “interaction” or “non-interaction”. The object-based and odour-based classifiers were trained on 28,586 and 32,872 frames of target “interaction” behavior, respectively. Prior to manual annotation, trimmed videos and filtered pose-estimation data was imported, then a scale factor was used to normalize variable camera filming heights to a known metric distance (experimental apparatus, dimensions = 60cm x 60cm). Additionally, each stimuli position was assigned a region-of-interest that was centered at each Velcro stimuli attachment point, with a defined radius extending ~2cm beyond the edge of stimuli. In total, 273 features were extracted from tracking data, where 251 features capture spatiotemporal relationships between points-of-interest, and 12 features capture ROI-related movement. We slightly deviated from the standard SimBA feature engineering approach by removing ROI-related features called “zone_cumulative_percent” and “zone_cumulative_time”. These features increase the prediction probability of a true class based on animal’s preferentially spending time in a defined ROI. While these features may be useful for predicting behaviors that only include in specific regions (e.g., rat dams retrieving pups from a nest), inclusion of these features in our project would bias predictions unequally between the six stimuli positions. For both the object and odor classifiers, the behavioral features most heavily weighted for model predictions include distance to stimuli, nose movements, region-of-interest, and spatial dynamics between points-of-interest (Supplementary Fig 2). Feature importance clusters were created by extracting the 40 most important features from SimBA, then splitting features based on the following criteria: 1) features related to the distance to stimuli “distance to stimuli”; 2) features related to nose movements (e.g., Nose_movement_M1_sum_6) were clustered to “nose movements”; 3) features related to a subjects’ nose key point being located within a defined ROI surrounding stimuli were clustered to “region-of-interest”; 4) remaining features were clustered to a common “spatial dynamics between points-of-interest”.

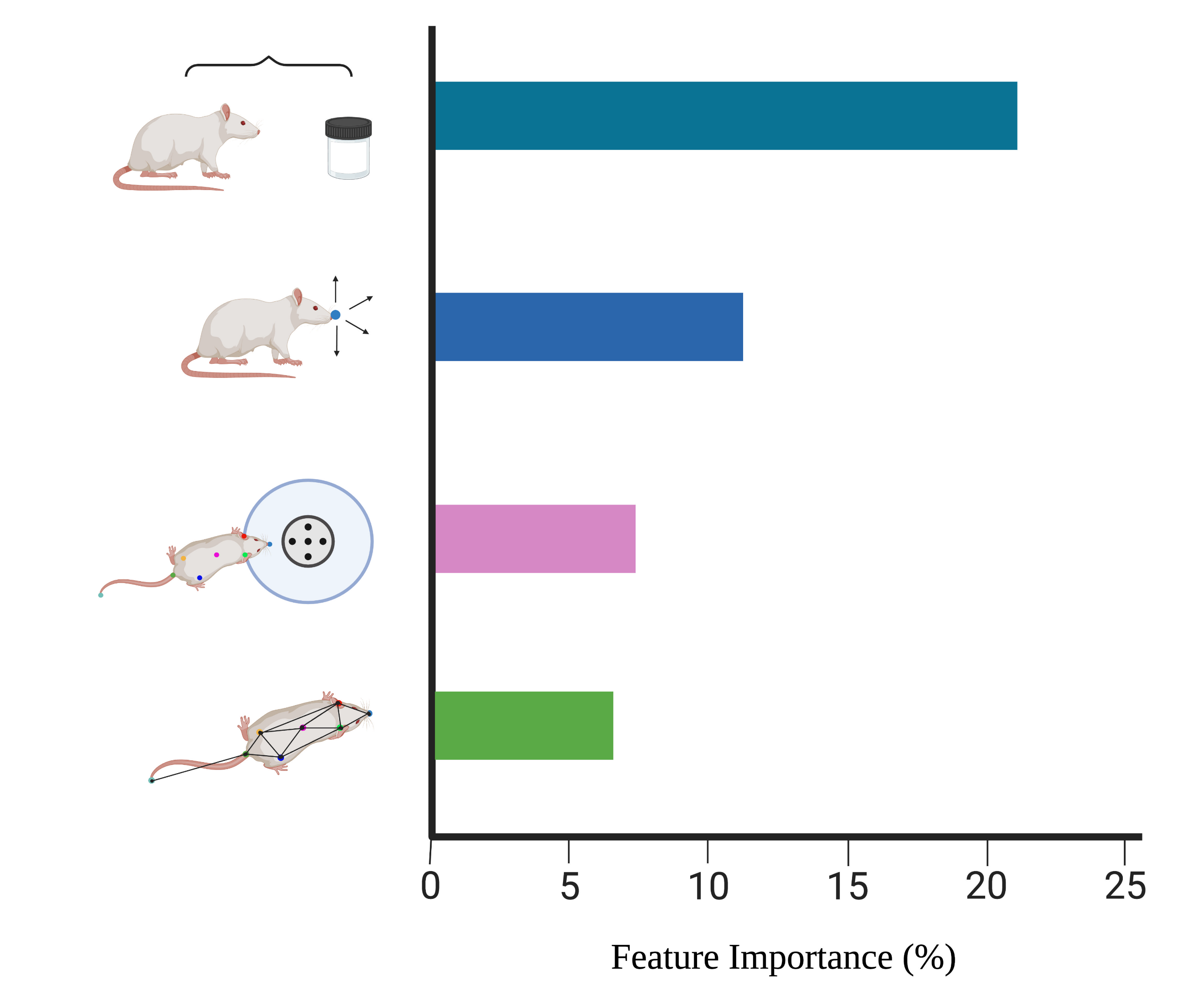

**Supplementary Fig 3.** Visualization of the relative feature importance of the four features clusters. In short, the 40 most important features were systematically categorized into distinct clusters, then we summed the feature importance’s of individual features within each cluster. The raw features importance log is included under “assessment + logs” for each classifier within our GitHub repository. This figure was created using BioRender.

For object-based videos, “interaction” was operationally defined as frames where the nose is within 2cm of the object, while looking at and/or chewing the stimuli for a duration greater than 50ms. For odor-based videos, “interaction” was operationally defined as frames where the rats’ nose is within 2cm, and within 2cm from the top of the odor jar, looking at and/or chewing the stimuli for a duration greater than 50ms. Classifiers were built using the following hyperparameter set: n_estimators = 200, RF_criterion = entropy, RF_max_features = sqrt, RF_min_sample leaf = 2 (Supplementary Fig 3).

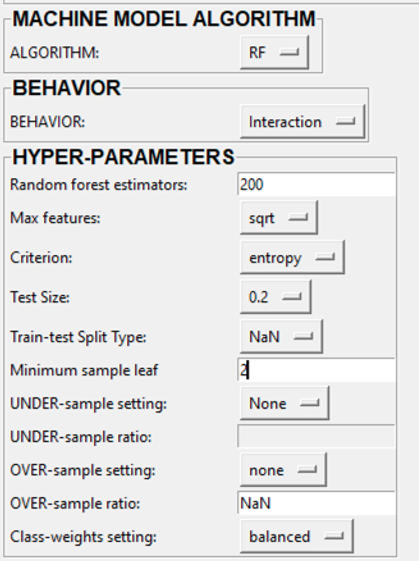

**Supplementary Fig 4.** Model hyperparameters used for classifier training. A meta-data csv file is included under “assessment + logs” for each classifier within our GitHub repository.

Previous studies have shown that creating a balanced dataset by using the model hyperparameters of “random under sampling” or “random over sampling” lead to better classifier performance; however, we found that using these features dramatically decreased classifier performance and lead to equal classifier predictions across the data frame. Therefore, we chose to not use these hyperparameters for analysis, and accounted for the unbalanced dataset by setting a relatively low discrimination threshold. For both classifiers, a discrimination threshold of 0.35 and a minimum bout duration of 50ms was used (Supplementary Fig 4).

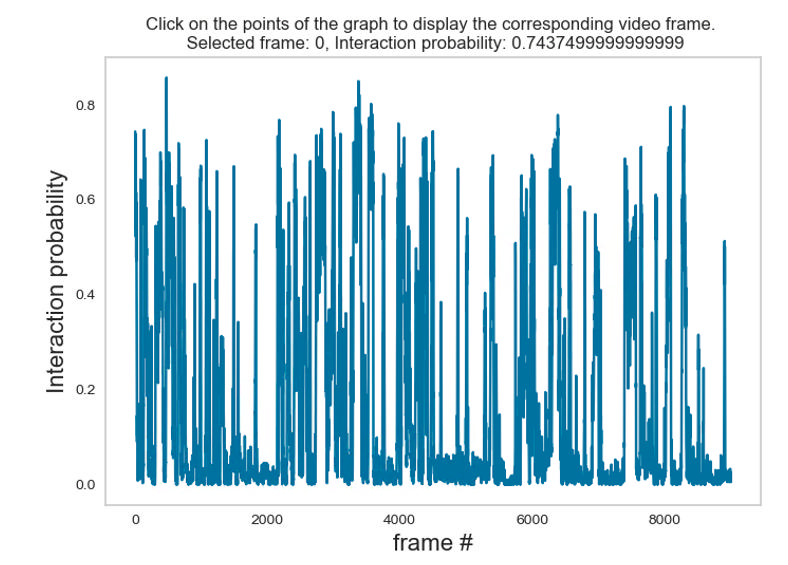

**Supplementary Fig 5.** Representative plot of classifier predictions across a complete video (9000 frames, 5 min video). We chose a discrimination threshold of 0.35 as it corresponds to the middle segment of obvious probability spikes and excludes the majority of noise below 0.2.

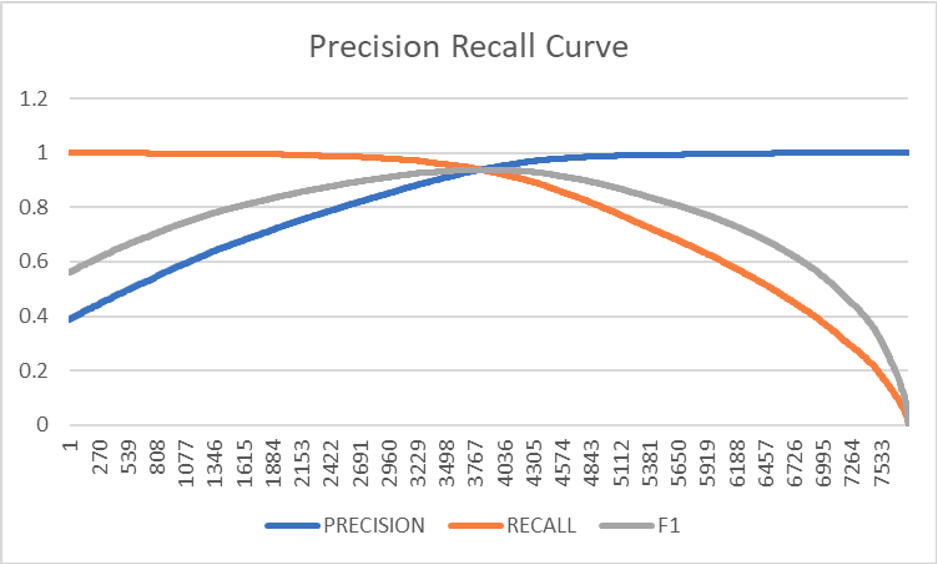

**Supplementary Fig 6.** Precision recall curve visualizing changes in precision, recall, and F1 with classifier training. Raw data is included under “assessment + logs” for each classifier within our GitHub repository. A detailed explanation of precision, recall, and F1 can be found below.

Recall, precision, and by extension the F1 score are calculated from the entries of a confusion matrix. A confusion matrix tells us, given a set of observations belonging to at least 2 different classes and a classifier that attempts to label each, how many and what type of errors were made. The diagonal of the confusion matrix is the correct observations, the off diagonal are the errors. For a binary classifier, we are generally focused on one class over the other, thus the metrics we derive are chosen to represent how we did for the most important class. In our case 'interaction' is the class we care about. In quantifying how our classifier for 'interaction' did, we calculate the recall and precision. Recall is the proportion of all the possible 'interaction' observations that our classifier predicted correctly. That is, the number of True Positives (TP) divided by the total number of 'interaction' observations (note the maximum number of True Positives is all the 'interaction' observations, in which case the recall equals 1, so a classifier that always predicts interaction will have perfect recall). Now there are many other metrics that could be computed, but the next most natural is the precision. Precision is the proportion of predicted 'interaction' observations that were actual 'interactions'. Or mathematically, the number of True Positives divided by the total number of times our classifier predicted 'interaction' (note it's not so easy to get perfect precision). Now we have 2 perfectly good numbers that quantify how our classifier did, the proportion of overall 'interactions' that were recovered (recall) and the proportion of times our classifier predicted 'interaction' and was correct (precision). It's not clear which is more important, so we combine the two (Supplementary Fig 5). We define the F1 score to be the harmonic mean of recall and precision. Why harmonic mean? We want an average of some kind, and the harmonic mean is the smallest of the 3 Pythagorean means (arithmetic mean, geometric mean and harmonic mean). So, to have a high F1 score you must have high precision and recall, either one will drag the F1 score down non-linearly.

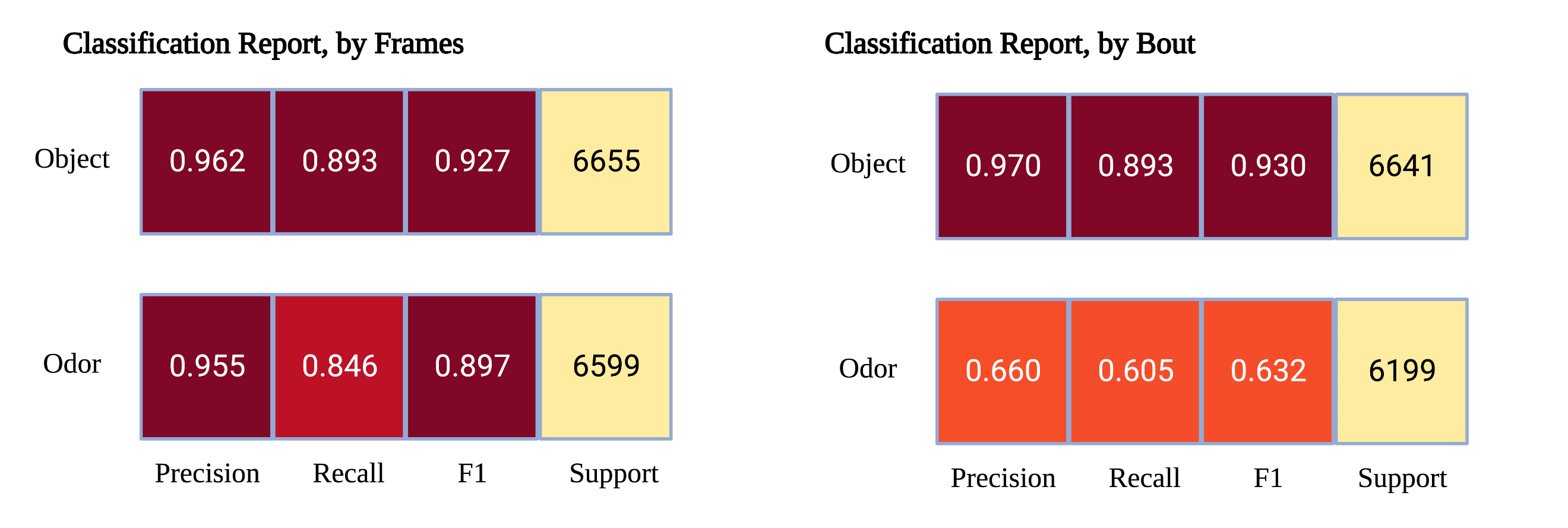

**Supplementary Fig 7**. Classification reports for object and odor (right) based analysis. Metrics are calculated by splitting the training dataset by-frames (left) and by-bout (right). This figure was created using BioRender.

We assessed model performance in two ways, both of which are integrated in the SimBA GUI (Supplemental Fig 6). First, we generated performance metrics (precision, recall, F1) by randomly splitting the aggregate training set (all human-annotated frames from all videos within the project) into 80% training frames and 20% test frames. Said differently, for a given behavioral video, a fraction of interaction-containing frames was used for model training, then a smaller fraction of frames was used for testing if the model can accurately predict if rat-stimulus interaction occurs in each test frame. As shown below, we found that both the object and odour classifiers generated excellent performance metrics when assessed in this manner. However, a fundamental problem with this assessment method is that for a given interaction bout, there may be both test and training frames, so the model is predicting interaction between two known sub-bouts of interaction (visualized- 1 = known interaction, test = test frame that the model must make a prediction on: 1-1-1-1-1-test-1-1-1-1). Therefore, to assess performance without the confound of intra-bout test frames, we segregated the aggregate training into interaction bouts, then split the segregated training set into 80% training bouts and 20% test bouts. We found that the performance of the object classifier changed marginally with this change, but performance metrics for the odor classifier significantly decreased when assessed in this manner. While we content that assessing classifier performance by-bout is a more conservative and representative method, an important caveat is that classifier performance on a completely model-naïve video is not assessed by either of these methods. This is important to consider because researchers will typically implement this analysis method to automatically quantify behavior for a large dataset, where only a fraction of this dataset is used for training. We did not include a by-video classifier analysis as this is not integrated into SimBA, but we contend that future research and software development should implement this performance assessment method to capture the accuracy of classifier predictions most accurately on model naïve behavioral videos.
